# Gapless assembly of maize chromosomes using long read technologies

**DOI:** 10.1101/2020.01.14.906230

**Authors:** Jianing Liu, Arun S. Seetharam, Kapeel Chougule, Shujun Ou, Kyle W. Swentowsky, Jonathan I. Gent, Victor Llaca, Margaret R. Woodhouse, Nancy Manchanda, Gernot G. Presting, David A. Kudrna, Magdy Alabady, Candice N. Hirsch, Kevin A. Fengler, Doreen Ware, Todd P. Michael, Matthew B. Hufford, R. Kelly Dawe

## Abstract

Creating gapless telomere-to-telomere assemblies of complex genomes is one of the ultimate challenges in genomics. We used long read technologies and an optical map based approach to produce a maize genome assembly composed of only 63 contigs. The B73-Ab10 genome includes gapless assemblies of chromosome 3 (236 Mb) and chromosome 9 (162 Mb), multiple highly repetitive centromeres and heterochromatic knobs, and 53 Mb of the Ab10 meiotic drive haplotype.

---

Maize is a classic genetic model, known for its excellent chromosome cytology and rich history of transposon research^1^. Transposons make up the majority of the maize genome^2^, and their accumulation over millions of years has driven genes far apart from each other and separated genes from their regulatory sequences^3^. There are also large inversions and other structural variations that contribute to fitness^4,5^ and significant variation in genome size caused by tandem repeat arrays ^6^. Understanding this remarkable structural diversity is important for the continued improvement of maize, but the high repeat content has impeded progress^2,5^. Here we describe an automated assembly merging approach that yields gapless maize chromosomes and dramatically improves contiguity throughout the genome, including centromere and knob regions.

The most challenging genomic regions to assemble are tandem repeat arrays that exceed the read length of the current sequencing technologies. In most eukaryotes, these arrays are enriched in centromeres and ribosomal DNA (rDNA). Maize contains a centromeric repeat of 156 bp^7^, an 45S rDNA repeat of 9349 bp, and a 5S rDNA repeat of 341 bp. In addition, maize contains two abundant classes of knob repeats that are found on chromosome arms, the major knob180 repeat (180 bp)^8^ and the minor TR-1 repeat (360 bp)^9^. Knob repeats occur in arrays that extend into the tens of megabases and present a significant barrier to full genome assembly. In most maize lines, knobs appear as inert heterochromatic bulges^8^, but in lines with a meiotic drive system on Abnormal chromosome 10 (Ab10) they have centromere-like properties and are preferentially segregated to progeny^10^. Ab10 is considerably longer than chromosome 10 and contains two inversions^11^, three knobs, and long spans of uncharacterized DNA that include a cluster of *Kinesin driver* (*Kindr*) genes required for meiotic drive^9^. Meiotic drive systems have been documented in many organisms and often lie within large inversions that contain novel repeat arrays^12^, yet no meiotic drive haplotype has been fully sequenced and assembled.

A new maize inbred, B73-Ab10, was created by backcrossing a line containing Ab10 to the B73 inbred six times and selfing it an additional five times (BC_6_F_5_). We used DNA from this line to prepare an optical map with the Bionano Saphyr system and sequenced it to high coverage using both PacBio and Nanopore technologies. We then implemented a genome assembly workflow based around the optical map (Suppl. Figure 1). Briefly, the PacBio data were assembled using Canu^13^, the Nanopore data assembled using miniasm^14^ and the two independent assemblies merged with miniasm and integrated with the optical map as hybrid scaffolds. Hybrid scaffolds were then used to guide further gap closing and create a pseudomolecule assembly (Figure 1A) where the superior PacBio assembly provides the core and Nanopore contigs help to seal the gaps. Our approach of one-step contig merging and error correction using optical maps as a reference differs from other methods that rely on local assemblies to fill gaps and correct errors^15,16^ and performs well in difficult repeat-rich regions (Figure 1B). Alignments of the optical map to the independent assemblies^17^ and standard genome completeness measures demonstrate that the approach is highly accurate (Suppl. Table 1 and 2).

**Figure 1.**
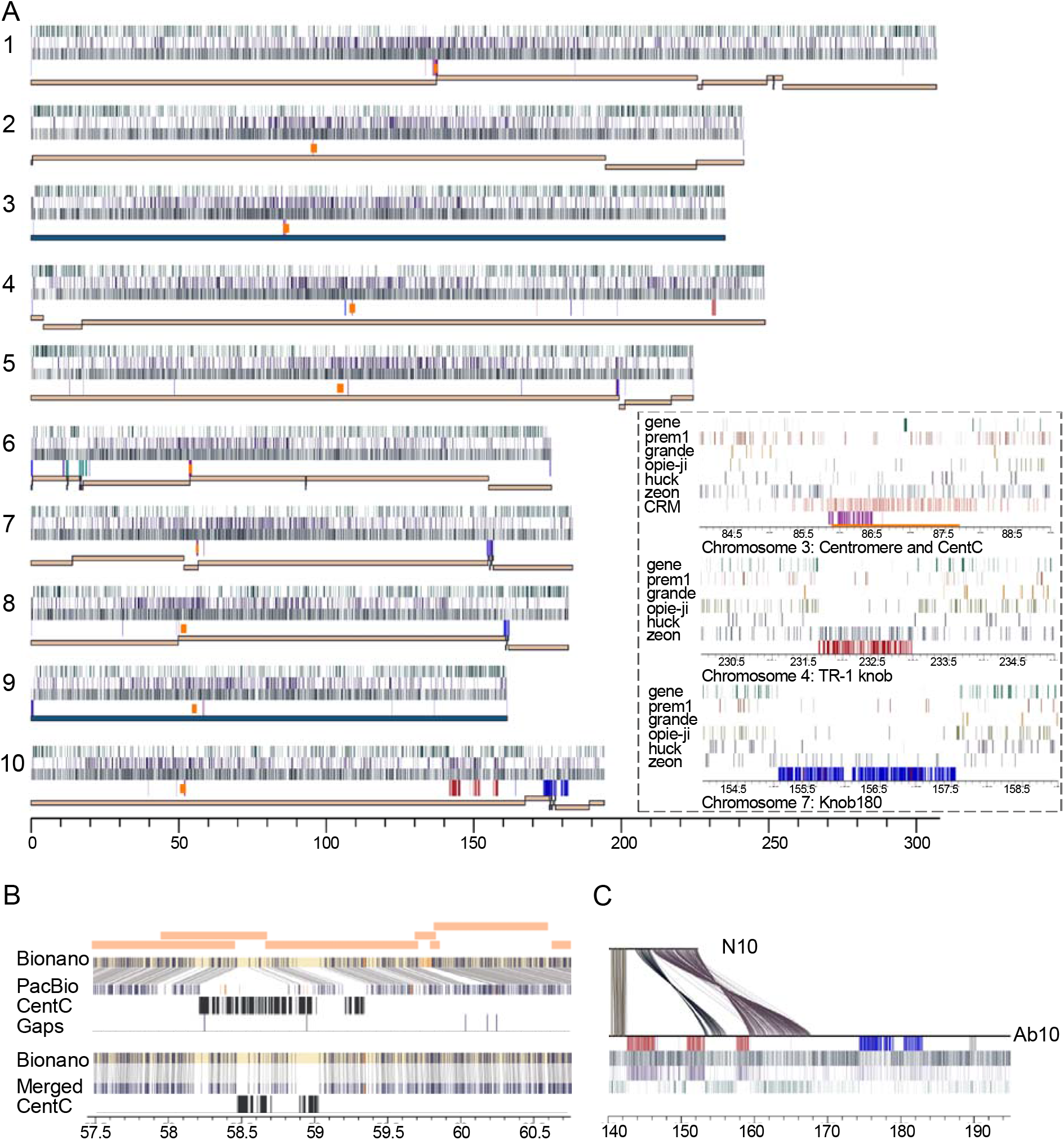
Assembly of the B73-Ab10 genome. **A**) Whole-genome view. For each chromosome, the top to bottom tracks are: gen density, cinful-zeon retrotransposon density, *Gypsy* superfamily retrotransposon density in 10 Kb sliding windows, repeat locatio (knob180 in blue, TR-1 in red, 45S rDNA in teal, CentC in magenta), and the distribution of gapless contigs. CENH3 ChIP-seq peaks identifying centromeres are marked by orange rectangles. The inset shows the centromere on chromosome 3, TR-1-rich knob on chromosome 4, and knob180-rich knob on chromosome 7. The five most common retroelement families are shown for each panel, along with Centromeric Retrotransposons (CRM) for the centromere. **B**) The impact of assembly merging over a CentC-rich region on chromosome 9. Seven contigs (orange, above) from the PacBio assembly were originally misassembled, a can be seen in the alignment to the Bionano map (connecting lines show matching sites). CentC tracts and gaps are annotated. Assembly merging corrected the output, leaving an 11 kb gap that was filled with nanopore reads. **C**) Sequence alignment between normal chromosome 10 from B73 (N10) (140Mb-152Mb) and Ab10 (140Mb-195Mb) from B73-Ab10. Annotation is as in A, with *Kindr* genes marked with black bars in the top track. Links show homologous regions larger than 500bp.

The final assembly has a contig N50 of 162 Mb (Table 1), which far exceeds the contiguity of any prior maize genome assembly^2,5^. Of particular note is the complete 236 Mb assembly of chromosome 3, which was assembled gaplessly without manual intervention – a first for any chromosome from a large complex genome. While the human X-chromosome was also assembled gaplessly^18^, this outcome required extensive manual inspection and correction. The entire B73-Ab10 genome is represented by 63 contigs where 90% are longer than 20.4 Mb (the N90). In addition to the expected gaps in repeat arrays, there were two gaps associated with residual heterozygosity on chromosome 9. Regions of heterozygosity reduce effective coverage and lead to assembly chimeras that are broken during hybrid scaffolding. We filled these heterozygosity-associated breaks by choosing the dominant Bionano path and performing local assemblies over the gaps. Nanopore reads were also used to span a gap within a CentC array to complete the chromosome 9 telomere-to-telomere assembly. Aside from these manual interventions, some efforts to manually improve within-knob assemblies, and a correction to the *Kindr* gene complex region of Ab10, the assembly was automated.

**Table 1.** Assembly metrics of the B73-Ab10 genome.

|  | **Contigs** |  |  | **Pseudomolecules** |  |  |
| --- | --- | --- | --- | --- | --- | --- |
|  | N50 (Mb) | N90 (Mb) | Max Size (Mb) | Contig Number | Total Length (Mb) | Gap<sup>a</sup> Length (Mb) |
| **Nanopore** | 2.0 | 0.5 | 8.3 | 1673 | 2161.1 | 93.2 |
| **PacBio** | 41.2 | 7.1 | 156.3 | 216 | 2162.7 | 2.6 |
| **Merged** | 162.0 | 20.4 | 235.9 | 63 | 2162.8 | 1.3 |

<sup>a</sup> Gaps longer than 10 Ns.

Seven of the ten functional centromeres as defined by ChIP-seq of CENP-A/CENH3^7^ were assembled without gaps (Suppl. Table 3). These include centromere 3, where CENH3 localizes over a 771-kb CentC array interspersed with transposons. Alignment of partial BAC-ased assemblies of B73 centromeres showed excellent agreement overall (Suppl. Figure 2). Prior maize assemblies have succeeded in obtaining only small fragments of knob repeat arrays. In contrast, a knob180-rich knob on chromosome 9 (850 kb), a TR-1-rich knob on chromosome 4 (1.3 Mb) and three TR-1-rich knobs (4.2 Mb, 2.6 Mb, and 2.1 Mb) on Ab10 were fully assembled in the B73-Ab10 assembly. The internal structure reveals that knobs, like centromeres^2,7^, often contain more transposons than tandem repeats (Figure 1C). Centromeric Retrotransposons target areas with CENP-A/CENH3^2,7^ and occupy on average 31.9% of functional centromeres, including within CentC arrays (Figure 1A and Suppl. Table 3 and 5). The new knob assemblies reveal that the Cinful-Zeon family of *Gypsy* elements^19^ preferentially target knobs. Cinful-Zeon elements occupy 27.0% of the assembled TR-1-rich knobs and 8.2% of the knob180-rich knobs, but only 3.8% percent of CentC arrays (Figure 1A and Suppl. Table 5 and 6). While Centromeric Retrotransposons are centromere-specific, Cinful-Zeon elements are also abundant in other heterochromatic regions throughout the genome (Figure 1A).

In addition to revealing the internal structure of knobs, the data provide the first complete view of the Ab10 haplotype that provides the selective force for the accumulation and maintenance of knobs^10^. The meiotic drive haplotype on Ab10 contains three fully assembled TR-1 knobs, a much larger knob180 knob that is not assembled, and two large inversions (4.4 and 8.3 Mb) that are homologous to normal chromosome 10 (Figure 1C). These major structural differences help to explain why recombination between the Ab10 haplotype and normal chromosome 10 is suppressed^20^. Ab10 also contains 22.4 Mb of novel sequence with no synteny to other regions of the maize genome or related grass genomes. Within this domain is the complete cluster of nine *Kindr* genes that are integral components of the drive system^10^, as well as hundreds of other expressed genes, many of which have only one exon or overlap with transposons and are likely non-functional (Suppl. Table 7). Additional meiotic drive functions associated with the movement of knobs at meiosis and their delivery to egg cells^21^ remain to be identified in this newly discovered sequence.

Gapless genome assemblies remove all uncertainty about the order, spacing and orientation of genes and their regulators. We have shown that this can be achieved using long reads and well-known assembly algorithms, with significant improvements in contiguity obtained by integrating independent assemblies around an optical map scaffold. Given that most contigs end in telomeres, centromeres or knobs, we presume that virtually all of the genes and associated regulatory information are represented in this genome assembly. The assembly merging pipeline also revealed the internal structure of repetitive domains that were previously known only by cytological techniques, thereby opening these regions to annotation and future epigenomic profiling. Similar results should be achievable other complex genomes, although higher sequence coverage, longer reads, and/or additional scaffolding information may be needed for species with polyploidy or higher levels of heterozygosity.

## ONLINE METHODS

### Germplasm, data, and code availability

The B73-Ab10 inbred can be obtained as PI 690316 at the Germplasm Resources Information Network (GRIN), Ames, Iowa. All genomic sequence and Bionano data can be obtained at the NCBI SRA under Bioproject PRJEB35367. The RNA-seq data is deposited in EBI (Accession number E-MTAB-8641). The code used in this study is available at the GitHub repository https://github.com/dawelab/Ab10-Assembly.

### PacBio assembly

High molecular weight DNA was extracted from young leaves using the protocol of Doyle and Doyle^1^ with minor modifications. Young maize leaves flash frozen at −80°C were ground to a fine powder in liquid N2 followed by very gentle extraction in CTAB buffer (that included proteinase K, PVP-40 and beta-mercaptoethanol) for 1 hr at 50C. After centrifugation, the supernatant was gently extracted twice with 24:1 chloroform:iso-amyl alcohol. The upper phase was adjusted to 1/10^th^ volume with 3M KAc, gently mixed, and DNA precipitated with iso-propanol. DNA was collected by centrifugation, washed with 70% EtOH, air dried for 20 min and dissolved thoroughly in 1x TE at room temperature.

Sequencing libraries were constructed following PacBio’s template prep protocols (Procedure & Checklist – Preparing gDNA Libraries Using the SMRTbell Express Template Preparation Kit 2.0, PN 101-693-800 Version 01) for the Express Template Prep Kit 2.0 (Cat# 100-939-900) and sequenced using Sequel SMRTLink V5.1 and Sequel binding and sequencing chemistry v2.1. The longest 50X out of 62X PacBio raw sequences were error-corrected using falcon_kit pipeline v0.7^2^ without repeat masking by TANmask and REPmask (-e 0.75 -l 3000 --min_cov 2 --max_n_read 200). The error-corrected reads (43X, N50= 22.3 Kb) were then trimmed and assembled with Canu^3^ (v1.8) with the following parameters: correctedErrorRate=0.065 corMhapSensitivity=normal ovlMerThreshold=500 utgOvlMerThreshold=150. The accuracy of the Canu-generated contigs was increased by aligning the raw PacBio reads to the assembly using pbmm2 (v1.2.0) from pb-assembly^2^ and running the PacBio consensus algorithm tool Arrow (v2.3.3) (https://github.com/PacificBiosciences/GenomicConsensus) with default parameters to generate sequenced polished contigs. The contig assembly was further polished using 73X PE150 Illumina sequence by first aligning the reads to the Arrow polished assembly using minimap2^4^, followed by running the assembly tool Pilon^5^ (v1.22) to correct individual base errors and small indels using the following parameters: --fix bases --minmq 30.

### Nanopore assembly

Two different DNA extraction methods were used to generate high molecular weight (HMW) DNA for Oxford Nanopore (ONT) sequencing. CTAB DNA was prepared as described above for the PacBio assembly. Nuclear DNA was prepared using the protocol of Luo and Wing^6^ with minor modifications. Young leaves flash frozen at −80°C were ground with liquid nitrogen and incubated with NIB buffer (10 mM Tris-HCL, PH8.0, 10mM EDTA PH8.0, 100mM KCL, 0.5 M sucrose, 4 mM spermidine, 1 mM spermine) on ice for 15 min. After filtration through miracloth, Triton X-100 (Sigma) was added to tubes at a 1:20 ratio, placed on ice for 15 minutes, and centrifuged to collect nuclei. Nuclei were washed with NIB buffer (containing Triton X-100) and re-suspended in 40 ml of the same buffer and centrifuged again. After removal of all liquid, 10 ml of Qiagen G2 buffer was added followed by gentle resuspension of nuclei; then 30 ml G2 buffer with RNase A (to a final concentration of 50 mg/ml) was added. Tubes were incubated at 37°C for 30 min. Proteinase K (Invitrogen), 30 mg, was added and incubated at 50 C for 2 hr followed by centrifugation for 15 min at 8000 rpm, at 4°C, and the liquid gently poured into a new tube. After gentle extraction with Chloroform:isoamyl alcohol (24:1), DNA was precipitated with two thirds volume iso-propanol. The DNA pellet was washed with 70% EtOH, air dried for 20 min and dissolved in TE at room temperature.

DNA from both the CTAB and nuclear prep was used to generate either a rapid (SQK-RAD004) or one dimensional (1d; SQK-LSK109) sequencing library for ONT. The resulting libraries were run on either a MinION or GridION sequencer running for 48 hrs. All bases were called on the GridION using Guppy (v2.0), and the resulting fastq files were used for genome assembly. A total of 121 Gb (~50x) of ONT sequence was generated over 27 MinION flowcells. The data were filtered for reads >10 Kb using seqtk (https://github.com/lh3/seqtk), resulting in an estimated 30x coverage of the maize genome. The resulting reads were aligned (overlap) with minimap2 (v2.13;-x ava-ont -t 64)^4^ and an assembly graph (layout) was generated with miniasm (v0.3; -f <reads= <overlaps=)^7^. The resulting graph was inspected using Bandage^8^. A consensus genome assembly was generated by mapping reads >10 Kb to the assembly with minimap2, and then running racon (v1.3.1)^9^; the consensus process was repeated three times. The contig assembly was further polished using 73X PE150 Illumina sequence by first aligning the reads to the consensus assembly using minimap2^7^ followed by running the assembly tool pilon (v1.18)^5^ two times using 73X PE150 Illumina sequence.

### Optical map assembly

Ultra high molecular weight DNA was isolated from maize seedlings using a modified version of the Bionano Genomics Plant Tissue DNA Isolation Base protocol. Approximately 0.5 g of healthy aerial tissue was collected from young B73-Ab10 etiolated seedlings grown in soil-free conditions for 2 weeks. The leaves were treated with a 2% formaldehyde Bionano fixing solution, washed, chopped and homogenized using a Qiagen TissueRuptor in homogenization buffer. Free nuclei were pelleted at 2,000X g, washed, isolated by gradient centrifugation, and embedded in a low melting point agarose plug. The nuclei were lysed by treating with proteinase K and RNase A treatments as described previously^10^, and washed four times in Wash Buffer and five times in TE buffer. The purified high molecular weight nuclear DNA was recovered by melting the plug, digesting it with agarase and subjecting the resulting sample to drop dialysis against TE.

The Bionano Saphyr platform was used in combination with the Direct Label and Stain (DLS) process to generate chromosome-level sequence scaffolds^11^. Direct labeling was performed using the Direct Labeling and Staining Kit (Bionano Genomics, San Diego CA) according to the manufacturer’s protocol, except that one microgram of DNA was used and DNA Stain was added to a final concentration of 1 microliter per 0.1 microgram of final DNA. The labeled sample was loaded into a Saphyr chip and molecules separated, imaged and digitized using a Saphyr and Compute server. Data visualization, map assembly and hybrid scaffold construction were performed using Bionano Access (v1.3) and Bionano Solve (v3.4.0). A subset of 1,580,077 molecules with a minimum size of 150 Kb and combined length of 424,488 Mb were assembled without pre-assembly using the non-haplotype, no-CMPR-cut parameters without extend-split.

### Assembly merging and gap closing

We developed a pipeline to integrate independent contig assemblies and curate assembly errors using Bionano maps as an anchor. The pipeline consists of five steps: 1) conflict resolution, 2) assembly error curation, 3) contig merging, 4) hybrid assembly and contig overlap removal, and 5) manual curation and gap filling (Suppl. Figure 1). The first four steps were automated. A gapless chromosome 3 was generated upon contig merging in the third step, and the complete assembly of chromosome 9 required manual curation. While contig merging with miniasm can be applied to any two sequence assemblies, the availability of *de novo* assembled Bionano maps is necessary to perform conflict-cutting in step 1, contig error correction in step 2, and hybrid scaffolding in step 4 of the pipeline.

Step 1: Conflicts between the optical map and DNA sequence assemblies were resolved using Bionano Solve software (https://bionanogenomics.com/support-page/data-analysis-documentation/). Sequence assembly can occasionally connect two regions that share a repetitive sequence but do not belong together (making a chimeric contig). These appear as conflicts between bionano maps and sequence assemblies when they are aligned. Optical maps were aligned to *in silico* digested representations of the DNA sequence assemblies using RefAligner (v3.4.0) and conflicts identified with the AssignAlignType.pl script. Conflicts with chimeric a quality score higher than the default threshold were split using cut_conflicts.pl (using default parameters from optArguments_nonhaplotype_noES_DLE1_saphyr.xml) and a sequence file was produced with custom script cut_conflict_NGS.py. Removing chimeric joins increases the chance of complementary contig merging in Step 3.

Step 2: Assembly errors in the conflict-resolved PacBio contigs were identified and automatically curated with ONT contigs. In this step, PacBio and ONT contigs were aligned to rescaled optical maps and structural discrepancies detected using the structural variant calling pipeline from BionanoSolve (v3.4.0). Homozygous insertions and deletions with a confidence of at least 0.1 and size larger than 1 Kb were classified as true assembly errors in the PacBio contigs. On the condition that no structural discrepancies were found in the corresponding ONT contigs, the ONT contigs were used to replace the erroneous sequences in PacBio contigs using custom script SV_fix.py.

Step 3: ONT contigs were used to close gaps and improve contiguity of the PacBio contig assembly. ONT contigs were mapped to PacBio contigs with minimap2^4^ (v2.13; -k28 -w28 -A1 -B9 -O16,41 -E2,1 -z200 -g100000 -r100000 --max-chain-skip 100), and overlap regions merged using miniasm^7^ (v0.3; -1 -2 -r0 -e1 -n1 -h250000 -g100000 -o25000). This step creates PacBio/ONT hybrid contigs that are called unitigs. The unitigs were then combined with the remaining contigs from the PacBio backbone assembly to create a merged contig assembly. After this step a gapless chromosome 3 was generated (a region of heterozygosity from 164.5 to 166.2 Mb on chromosome 3 was automatically resolved). The merged contigs were then aligned to Bionano maps, where overlaps between adjacent contigs were detected and merged with minimap2 (v2.13) and miniasm (v0.3) using the custom script Overlap_merge.py. This step only identifies large overlaps (roughly >200 Kb) that can be detected at the level of *de novo* Bionano label alignment. Identifying all overlaps, including smaller overlaps, requires hybrid scaffolding with the optical map (Step 4). If proceeding to Step 4, overlap merging in Step 3 is optional.

Step 4: Bionano maps were integrated with the sequence contigs by hybrid scaffolding using the hybridScaffold.pl script from BionanoSolve (v3.4.0) with default parameters from optArguments_nonhaplotype_noES_DLE1_saphyr.xml. This step orders and orients sequence contigs and facilitates the resolution of remaining overlaps between contigs. As the optical maps are aligned and rescaled with the sequence maps repeatedly during hybrid scaffolding, more accurate overlaps between contigs are identified and annotated as 13N gaps. These overlaps were removed through contig merging with miniasm (v0.3), as described in Step 3. Due to the extreme repetitiveness in the 45S rDNA repeat region on chromosome 6, both the contig assemblies and hybrid scaffolding in this area are erroneous. Therefore, we left the contigs in the NOR un-merged and marked the incorrectness with 13N gaps.

Step 5: Manual curation was performed to correct assembly errors, close gaps in repetitive and heterozygous regions, and assemble telomeres.

#### Repeat assembly manual curation

In highly repetitive regions, erroneous read joins at the tips of contigs were not detected as conflicts or assembly errors in Steps 1 or 2 due to the limited resolution of Bionano alignment. In these regions, we trimmed and removed the unaligned regions to reveal eligible ends for overlap merging using miniasm (v0.3). These modifications extended the contiguity of repeat arrays at the edges of longer contigs. Contigs composed exclusively of knob and CentC repeats arrays lack pan-genome anchor markers and are not present in the pseudomolecules.

#### Chromosome 9 manual curation

Seven gaps, ranging from 2 Kb to 236 Kb, were present in the chromosome 9 assembly after hybrid scaffolding. Two large gaps of 236 Kb and 41 Kb were caused by heterozygosity (76.29-76.80 Mb), one 21 Kb gap was due to repetitiveness in a CentC array (58.43-58.67 Mb), and the remaining four gaps were smaller than 7 Kb (two of these were in the 843 Kb knob on the tip of 9S). The four small gaps were first filled by running three iterations of LR Gapcloser (Sep 24, 2018 commit)^12^ at default settings using PacBio error-corrected reads. To resolve the 236 Kb gap caused by heterozygosity, all contigs anchored to chromosome 9 were re-scaffolded using the longest chromosome 9 Bionano map as the sole anchor. This reduced the 236 Kb gap to 58 Kb. Local assemblies were run with Flye (v2.6)^13^ using ONT reads surrounding gaps to fill the remaining 58 Kb and 41 Kb gaps. Flye-assembled contigs were integrated with the flanking contigs by unitigging with miniasm (v0.3), and aligned to Bionano maps for inspection. An 8 Kb gap remained, which was filled with a single ONT read that spans it. The gap in the CentC array was filled by manually selecting two long ONT reads (>50 Kb) that spanned the gap, creating a consensus at the overlap and placing the resulting sequence in the gap.

#### Kindr complex manual curation

The assembly over the ~1 Mb tandem array of *Kindr* genes (each within an ~100 Kb repeat) was erroneous due to collapsing in the PacBio sequence contig and improper scaffolding. We manually selected the most contiguous ONT contig over this region, carried out hybrid scaffolding for the scaffold containing *Kindr*, placed an excluded contig in the correct area, and removed an overlap region through contig merging.

#### Telomere manual curation

Fifteen telomeres were assembled by extending the ends of scaffolds with the longest uniquely mapped ONT read that contained telomeric repeats TTTAGGG/CCCTAAA (>=1 Kb). The regions with newly assembled telomeres include 1L, 2L, 3S, 3L, 4S, 4L, 5L, 6L, 7S, 7L, 8S, 8L, 9S, 9L,10S.

The final scaffolds were polished with PacBio subreads using tools from pb-assembly^2^. Read alignment was performed with pbmm2 (v1.2.0) and polishing was executed with GCpp (v1.0.0) at default parameters. Scaffolds were further polished with 73X PE150 Illumina reads using Pilon (v1.23) with default parameters^5^.

### AGP construction

The pseudomolecules were constructed from the hybrid scaffolds using ALLMAPS (v0.8.12)^14^. Both pan-genome anchor markers^15^ and the IBM (Intermated B73 × Mo17) genetic map^16^ were used with equal weights for ordering and orienting the scaffolds. Pan-genome anchor markers were obtained from the CyVerse Data commons^17^ and processed to generate a bed file with 50bp upstream and downstream of B73 V3 coordinates. The extracted markers were mapped to a HiSat2 (v2.1.0)^29,30^ indexed assembly of B73-Ab10 by disabling splicing (--no-spliced-alignment) and forcing global alignment (--end-to-end). Very high read and reference gap open and extension penalties (--rdg 10000,10000 and --rfg 10000,10000) were also used to ensure full-length mapping of marker sequence. The final alignment was then filtered for mapping quality greater than 30 and tag XM:0 (unique mapping) to retain only high-quality, uniquely mapped marker sequences. The mapped markers were merged with the predicted distance information to generate a CSV input file for ALLMAPS. Only scaffolds with more than 20 uniquely mapped markers, with a maximum of 100 markers per scaffold, were used for pseudomolecule construction. The IBM genetic markers were downloaded from MaizeGDB (https://www.maizegdb.org/complete_map?id=887740)^18^ and were processed to generate a bed file similar to pan-genome markers. For the markers with coordinates, 50 bp flanking regions were extracted from the B73 v4 genome. For markers without coordinates, marker sequences were used as-is, and those missing both coordinates and sequences were discarded. Mapping of the markers was done similar to the method described above for the pan-genome anchor markers, with all uniquely mapped markers retained. The genetic distance information for these markers was converted to a CSV file before use in ALLMAPS. ALLMAPS was run with default options, and the pseudomolecules were finalized after inspecting the marker placement plot and the scaffold directions. Synteny dotplots were generated using the scaffolds as well as pseudomolecule assemblies against the B73 genome by following the ISUgenomics Bioinformatics Workbook (https://bioinformaticsworkbook.org/dataWrangling/genome-dotplots.html). Briefly, the repeats were masked using RepeatMasker (v4.0.9)^19^ and the Maize TE Consortium (MTEC) curated library (https://github.com/oushujun/MTEC)^20^. RepeatMasker was configured to use the NCBI engine (rmblastn)^21^ with a quick search option (-q) and GFF as a preferred output. The repeat-masked genomes were then aligned using minimap2^4^ (v2.2) and set to break at 5% divergence (-x asm5). The paf files were filtered to eliminate alignments less than 1 Kb and dotplots were generated using the R package dotPlotly (https://github.com/tpoorten/dotPlotly).

### RNA-seq

Ten tissues were sampled throughout development for evidence-based gene annotation including: primary root (1) and coleoptile (2) at six days after planting; base of the 10^th^ leaf (3), middle of the 10^th^ leaf (4), tip of the 10^th^ leaf (5) at the Vegetative 11 (V11) growth stage; meiotic tassel (6) and immature ear (7) at the V18 growth stage; anthers at the Reproductive 1 (R1) growth stage; endosperm (9) and embryo (10) at 16 days after pollination. For each tissue, two biological replicates were harvested and each biological replicate was made up of tissue from three individual plants. Endosperm and embryo tissues were harvested from 50 kernels per plant (150 total per biological replicate). Tissues 1-5 above were collected from greenhouse-grown plants and tissues 6-10 were from field-grown plants. Greenhouse-grown plants were planted in Metro-Mix300 (Sun Gro Horticulture) with no additional fertilizer and grown under greenhouse conditions (27°C/24°C day/night and 16h/8h light/dark) at the University of Minnesota Plant Growth Facilities. Field grown plants were planted at the Minnesota Agricultural Experiment Station located in Saint Paul, MN with 30-inch row spacing at ~52,000 plants per hectare. RNA was extracted using the Qiagen RNeasy plant mini kit following the manufacturer’s suggested protocol.

Total RNA samples were assayed by Bioanalyzer to determine RNA integrity and normalized in 25uL of nuclease-free water prior to library preparation. Sequencing libraries were prepared using KAPA’s Stranded mRNA-seq kit (#KK4821) according to the manufacturer’s instructions. The mRNA was enriched using oligo-dT beads, fragmented, and converted to double stranded cDNA using random hexamer priming and amplification. Libraries were pooled at equimolar ratios and sequenced on NextSeq 500 instruments using the PE75 protocol.

### Gene annotation

For evidence-based predictions, genome-guided transcript assemblies were generated from five different assemblers *viz.,* Trinity (v2.6.6)^21,22^, StringTie (v1.3.4a)^23^, Strawberry (v1.1.1)^24^, Cufflinks (v2.2.1)^23,25^ and Class2^23,25,26^, and the best set of transcripts were identified and annotated as genes using Mikado (v1.2.4)^27^. Briefly, the RNA-seq reads from each library were mapped to a STAR (v2.5.3a)^28^ indexed B73-Ab10 genome using a 2-pass mapping approach (the initial round of alignments provides splice information for the subsequent round of mapping reads). Default options were used for mapping with few post-processing options enabled (print all SAM format attributes --outSAMattributes All; downstream compatibility -- outSAMmapqUnique 10; and number of mis-matches --outFilterMismatchNmax 0). Individually mapped RNA-seq libraries were then pooled, sorted and indexed using SAMTools (v1.9)^29^, for use with the transcript assembly programs. For all genome-guided transcriptome assemblers, default options were used except, if it allowed minimum transcript length setting, it was set to 100 bp (Trinity using --min_contig_length 100, StringTie using -m 100 and Strawberry using -t 100), and if it allowed RNAseq strandedness, it was set to stranded (Trinity using -SS_lib_type FR, Cufflinks using --library-type fr-firststrand). For Trinity, maximum intron size was also set to 10000 (--genome_guided_max_intron 10000). All assemblers generated a GFF3 as the final output except for Trinity, for which assembled transcripts in fasta format were mapped back to the gmap (v2019-05-12) indexed genome to generate a GFF3 file (by setting the output format option -f to gff3_match_cdna). Portcullis (v1.1.2)^30^ was used to generate a high confidence set of splice junctions for the B73-Ab10 genome from the merged mapped reads. Mikado was configured to use all transcript assemblies (with strandedness marked as True for all except for Trinity, and with equal weights), portcullis generated splice sites and a plants.yaml scoring matrix. Preliminary transcripts prepared by Mikado, through merging all transcripts and removing the redundant copies, were processed using TransDecoder (v5.5.0)^31^ (to identify open reading frames) and blastx (v2.9.0)^21^ against SwissProt viridiplantae proteins (for identifying full-length transcripts). Default options were used for TransDecoder, and for blastx, maximum target sequences were set to 5 (-max_target_seqs 5) and output format to xml (-outfmt 5). These were provided as input for Mikado for picking and annotating the best transcripts for each locus. The obtained GFF3 file was used to extract transcripts and proteins using the gffread utility from the Cufflinks package.

Additional structural improvements for the Mikado generated transcripts were completed using the PASA (v2.3.3)^32^ genome annotation tool. The inputs for PASA included 2,019,896 maize EST derived from genbank, 83,087 Mikado transcripts, 69,163 B73 full length cDNA from genbank and 46,311 maize iso-seq transcripts from 11 developmental tissues that were filtered for intron retention^33^. PASA was run with default options, with a first step of aligning transcript evidence to the masked B73-Ab10 genome using GMAP (v.2018-07-04)^34^ and Blat (v.36)^35^. The full length cDNA and Iso-seq transcript ID’s were passed in a text file (-f FL.acc.list) during the PASA alignment step. Valid near perfect alignments with 95% identity were clustered based on genome mapping location and assembled into gene structures that included the maximal number of compatible transcript alignments. PASA assemblies were then compared with B73-Ab10 Mikado transcript models using default parameters. PASA updated the models, providing UTR extensions, novel and additional alternative isoforms. PASA generated models were passed through the MAKER-P (v3.0)^36^ annotation pipeline as model_gff along with all the transcript and protein sequences to obtain Annotation Edit Distance (AED)^37^ scores to assess the quality of annotations. Transposon element (TE) related genes were filtered using the TEsorter tool^21,38^, which uses the REXdb (viridiplantae_v3.0 + metazoa_v3) database of TEs. Finally the gene annotations were verified for translation errors using the EnsemblCompara pipeline^39^.

### BUSCO assessment

The gene space completeness of the B73-Ab10 genome assembly was assessed using the GenomeQC^40^ tool, which provides a summary of the number of complete, fragmented and missing Benchmarking Universal Single-Copy Orthologs (BUSCO) in the assembly. The Embryophyta database (embryophyta_odb9; consisting of 1440 conserved, single-copy plant genes) and the genome assembly in the fasta file format were provided as input to the tool to calculate the BUSCO metrics.

### TE annotation

The manually curated transposable element library (maizeTE11222019) derived from the Maize TE Consortium (MTEC; https://github.com/oushujun/MTEC) was used as the base TE library. Novel TEs of the maize Ab10 genome not included in the MTEC library were structurally identified using the EDTA pipeline (v1.6.5)^41^ with parameters “-species maize -curatedlib maizeTE11222019”. The MTEC library augmented with Ab10 specific TEs was used to annotate TE fragments using RepeatMasker. Coding sequences of the maize B73 v4 assembly were downloaded from MaizeGDB and used to remove gene sequences in the EDTA-generated TE library. Whole-genome TE annotations were generated using the EDTA augmented MTEC library (-anno 1). The LTR Assembly Index (LAI)^42^ scores of genome assemblies were calculated using LAI (beta3.2) within the LTR_retriever (v2.8)^43^ package with parameters “-iden 94.8550 -totLTR 76.34”.

### Centromere and repeat analyses

The overall accuracy of the centromere assemblies was assessed by aligning previous BAC-based B73 centromere assemblies^17^ to the B73-Ab10 genome using Bionano RefAligner (v3.4.0) with default parameters. Although the BAC-based assemblies do not traverse CentC arrays, there is excellent overall agreement in sequence and contiguity (Suppl. Figure 2).

Active centromere locations were determined by identifying the CENH3 ChIP-seq enriched regions in the final assembly using genomic reads as a control. The SE150 Illumina ChIP-seq reads were obtained from SRA (SRX2737618)^44^ and the 73X PE150 Illumina genomic reads were subsampled to 30X with seqtk (https://github.com/lh3/seqtk). Both the ChIP-seq reads and the genomic reads were trimmed with Trim Glore v0.4.5 (https://github.com/FelixKrueger/TrimGalore/) with default parameters and aligned to the final assembly with BWA-MEM v0.7.17^45^. Epic2^46^ was employed to call peaks with the CENH3 ChIP-seq alignment set as treatment, genomic read alignment as control, MAPQ (mapping quality) as 20, effective genome size as 0.8, bin size as 5000 and gap size as 0. The effective genome size of the final genome was calculated as the fraction of unique 150-mers over total 150-mers using Jellyfish v2.26^47^ (-m 150 -s 2193M -out-counter-len 1 -counter-len 1). The coordinates of active centromeres were identified as islands with a score above 250 and a fold change higher than 4.

The coordinates of repeat arrays were identified by blasting the knob180 and CentC consensus sequences^44^, a TR-1 consensus (TTCTTTATATTCCAACTTTTTAGCAACTGTATGGTGGAAAAAGGTGTCTTACAACCTTAACCTATGTTTGGACAGTTCTCTCGTGCAATTTGGCTAAATTTCCCATGGTCTTTATTTTATTTTGAGAAACGATGTGGTATAATGATGTGCGATGTTTTACTTGAGTGGACATAAACACCATTTAGGTATGCCTTGAATAGAGGGGATTATTGGAAACCTGGTATCACAAAAGGTCATTAGCTAGCCCAATAACGTCTTCATCCACTAGTTATACTCTAATACCCTCTAGTGTGAATACAATGCCCACAATATCATAGAAACGTCATTTGAGGTTTAAAAGGTGATCTATTGTTTTGAA), subtelomeric repeat (NCBI CL569186.1) and ribosomal DNA intergenic spacer sequences (NCBI AF013103.1) against the B73-Ab10 genome. Knobs were defined as repeat clusters (>=500 Kb) that are composed of at least 10% repeat consensus sequences (knob180 and TR-1) with no more than 100 Kb spacing between repeat units. This definition of knob180 knobs excludes the subtelomeric knob180 arrays. CentC arrays are defined as repeat clusters (>=100 Kb) that are composed of at least 10% CentC consensus sequences.

Non-overlapping repeat units were quantified in each repeat array with custom script repeat_analyses.py. Five major families of the long terminal repeat (LTR)-retrotransposons in knobs and CentC arrays were individually quantified with bedtools (v2.28.0)^48^. The *Opie-Ji* family includes *Opie, Ji, Ruda* and *Giepum* and the *Prem1* family is composed of *Prem1*, *Xilon, Diguus,* and *Tekay^49^*. Centromeric retrotransposons CRM1 and CRM2 were quantified together and annotated as CRM in active centromeric regions with bedtools (v2.28.0).

