## Supplementary Figure 1-2, Table 1-7 for "Gapless assembly of maize chromosomes using long read technologies"

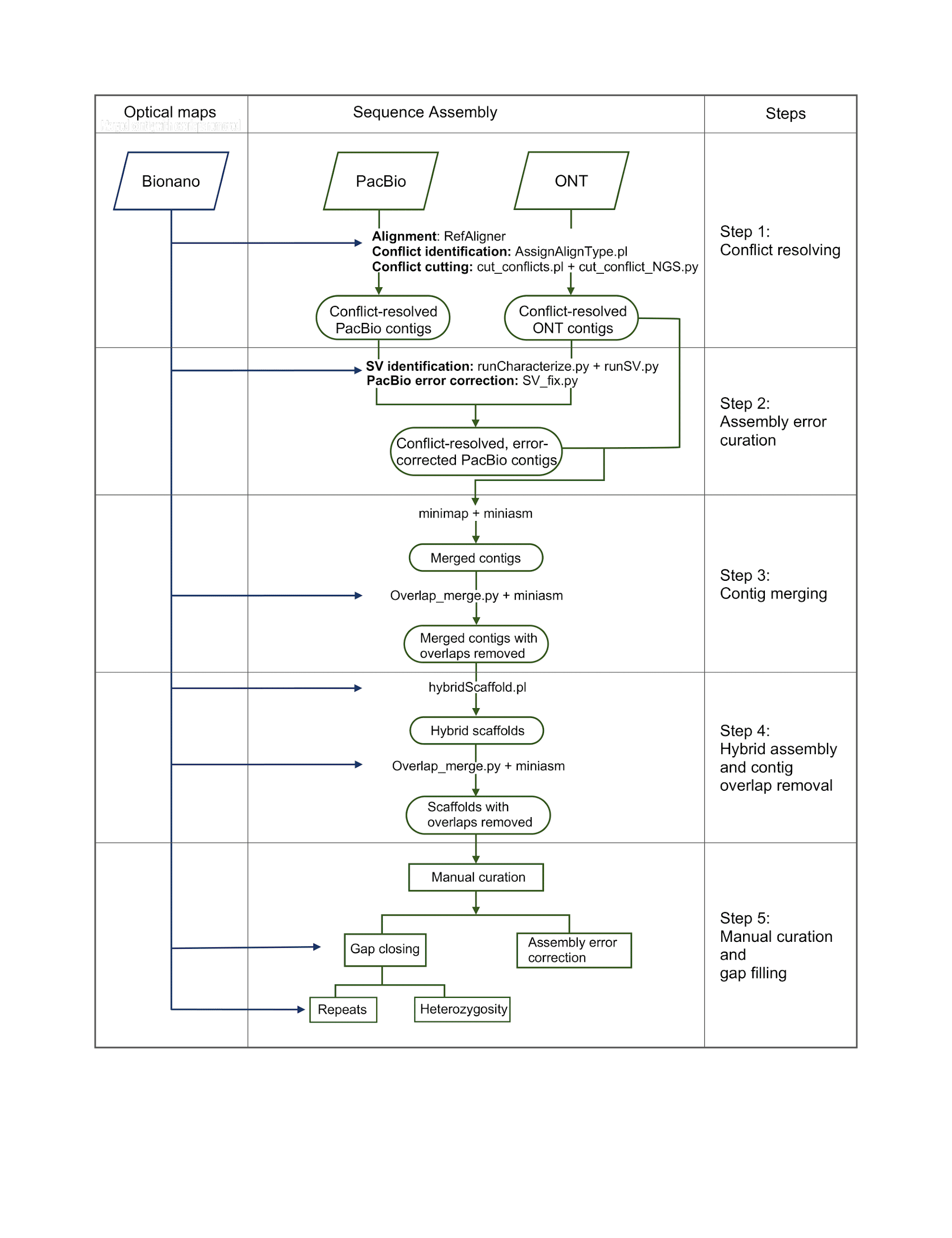

**Supplementary Figure 1.** Workflow for the B73-Ab10 assembly pipeline.

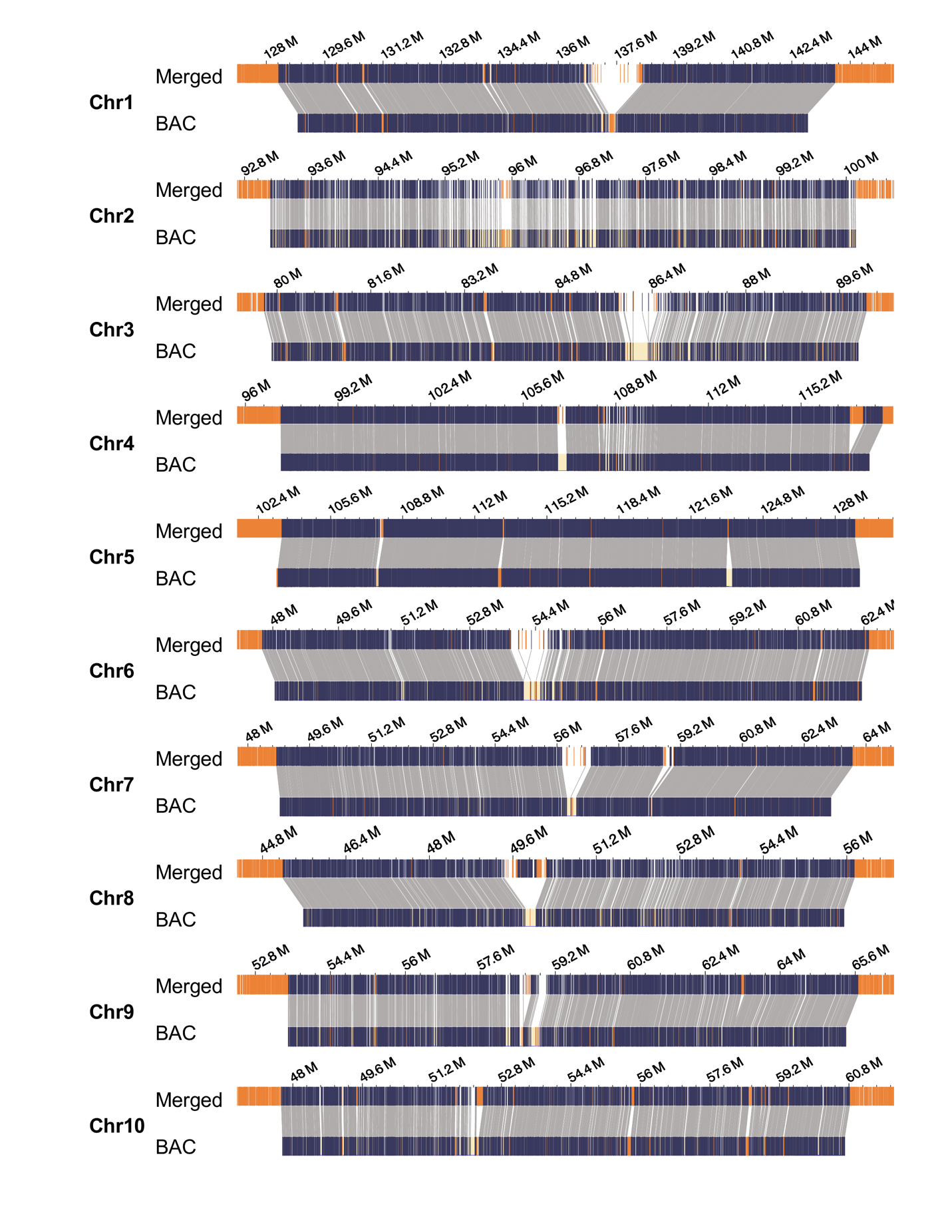

**Supplementary Figure 2.** The alignment of BAC-based assemblies of B73 centromeres to the merged assembly in optical map format. The connecting lines represent matching regions between the two assemblies.

|  | Metrics | Nanopore | PacBio | Merged |
| --- | --- | --- | --- | --- |
| Contig | N50 (Mb) / L50 | 2.0 / 325 | 41.2 / 16 | 162.0 / 6 |
|  | N60 (Mb) / L60 | 1.7 / 440 | 31.5 / 22 | 111.4 / 8 |
|  | N70 (Mb) / L70 | 1.3 / 584 | 23.8 / 30 | 88.8 / 10 |
|  | N80 (Mb) / L80 | 0.9 / 775 | 15.2 / 42 | 50.1 / 13 |
|  | N90 (Mb) / L90 | 0.6 / 1061 | 7.4 / 64 | 20.4 / 21 |
|  | Total contig number | 1912 | 1103 | 1016 |
|  | Total sequence (Mb) | 2120.5 | 2232.9 | 2241.4 |
|  | Contig number in Bionano scaffolds ^a^ | 1699 | 211 | 132 ^d^ |
|  | Contig sequence in scaffolds (Mb) ^a^ | 2072.5 | 2156.5 | 2174.2 |
| Scaffold | Number | 43 | 25 | 50 |
|  | N50 (Mb) | 125.7 | 162.9 | 161.8 ^e^ |
|  | Max (Mb) | 201.2 | 236.5 | 235.9 |
|  | Total length (Mb) | 2170.1 | 2162.2 | 2178.1 |
|  | Contig overlaps ^b^ | 929 | 114 | 12 |
|  | Gaps of known size ^c^ | 728 | 73 | 51 |
|  | Total known gap size (Mb) | 95.1 | 2.6 | 2.3 |
| Pseudo-  molecules | LTR Assembly Index (LAI) | 8.57 | 27.98 | 27.8 ^f^ |
|  | BUSCO (% Complete) | 90.7 | 95.6 | 95.8 |
|  | Total length (Mb) | 2161.1 | 2162.7 | 2162.8 |
|  | Contig overlaps ^b^ | 910 | 114 | 5 |
|  | Gaps of unknown size | 30 | 16 | 17 |
|  | Gaps of known size ^c^ | 728 | 73 | 31 |
|  | Total known gap size (Mb) | 93.2 | 2.6 | 1.3 |
|  | Assembled telomeres | 1 | 9 | 15 |

**Supplementary Table 1. Assembly statistics and gaps in B73-Ab10 assemblies.**

^a^ Contigs included in scaffolds are conflict-resolved contigs after hybrid assembly.

^b^ Contig overlaps are identified when the contigs are integrated with the optical map. The Bionano hybrid scaffolding software marks them with 13 Ns.

^c^ Gap sizes are estimated by Bionano maps during hybrid assembly.

^d^ Only 63 are included in pseudomolecules; the remaining 59 are anchored to small scaffolds (< 3Mb) containing CentC or knob arrays that lack genetic and pan-genome markers and could not be placed on chromosomes (Suppl. Table 4).

^e^ The N50 is smaller in Merged than PacBio due to the correction of the CentC array on chromosome 9 (Figure 1B).

^f^ LAI is lower in the Merged than PacBio due to a reduction in the total number of LTRs, presumably because sequence overlaps were removed.

**Supplementary Table 2. Accuracy of genome assemblies as assessed by comparison to Bionano maps.**

|  | Nanopore | PacBio | Merged |
| --- | --- | --- | --- |
| Contig misjoins / Conflict cuts | 425 | 18 | 1 |
| Number of collapsed repeats > 25 Kb | 705 | 56 | 3 |
| Sequence lost by collapse of repeats > 25 Kb (Mb) | 44.80 | 3.65 | 0.13 |
| Number of expanded repeats > 25 Kb | 22 | 13 | 10 |
| Sequence gained by expansion of repeats > 25 Kb (Mb) | 1.28 | 1.49 | 0.57 |

**Supplementary Table 3. Coordinates and composition of centromeres defined by CENH3 ChIP-seq in the B73-Ab10 assembly.**

| Chr | Start (bp) | End (bp) | Size (bp) | 100N^a^ | CentC (%) | CRM (%) | cinful-zeon (%) | grande (%) | huck (%) | opie-ji (%) | prem1 (%) |
| --- | --- | --- | --- | --- | --- | --- | --- | --- | --- | --- | --- |
| chr1 | 137,090,000 | 138,130,000 | 1,040,000 | Y | 54.8 | 41.1 | 0 | 0 | 0 | 0.9 | 0.1 |
| chr2 | 95,290,000 | 97,165,000 | 1,875,000 | N | 1.3 | 31.5 | 14 | 3.2 | 6.3 | 2.5 | 7.6 |
| chr3 | 85,880,000 | 87,705,000 | 1,825,000 | N | 14.7 | 37.2 | 8.8 | 0 | 4.8 | 2 | 4.6 |
| chr4 | 108,485,000 | 110,135,000 | 1,650,000 | N | 1 | 44.6 | 10.2 | 0.5 | 3.6 | 3.5 | 6.3 |
| chr5 | 104,255,000 | 106,220,000 | 1,965,000 | N | 0 | 16.3 | 19.2 | 0.8 | 2.2 | 1.5 | 8.1 |
| chr6 | 53,825,000 | 54,655,000 | 830,000 | Y | 49.1 | 41.4 | 0.8 | 0 | 1.5 | 0.2 | 0.0 |
| chr7 | 56,220,000 | 56,860,000 | 640,000 | Y | 30.8 | 54.5 | 0.3 | 0 | 0 | 0 | 0.0 |
| chr8 | 51,095,000 | 52,795,000 | 1,700,000 | N | 0 | 25.3 | 17.9 | 3.9 | 1.5 | 1.3 | 4.3 |
| chr9 | 54,840,000 | 56,275,000 | 1,435,000 | N | 0 | 5.1 | 18.8 | 6.4 | 2.8 | 2.2 | 11.1 |
| chr10 | 50,845,000 | 52,505,000 | 1,660,000 | N | 9.3 | 22.3 | 13.1 | 1.8 | 3.2 | 5.2 | 7.1 |

^a^ Gaps marked by 100 Ns are of unknown size.

**Supplementary Table 4. Repetitive components in B73-Ab10 assemblies.**

|  | Repeat Type | Nanopore | PacBio | Merged |
| --- | --- | --- | --- | --- |
| Scaffold | knob180 (bp) | 3,962,210 | 9,893,818 | 16,861,684 |
|  | TR-1 (bp) | 2,324,807 | 4,984,957 | 7,253,994 |
|  | CentC (bp) | 708,654 | 3,233,366 | 2,881,960 |
|  | rDNA intergenic spacer (bp) | 185,738 | 789,045 | 755,420 |
|  | Subtelomere (bp) | 99,243 | 504,890 | 563,144 |
| Pseudo-molecules | knob180 (bp) | 3,469,793 | 9,894,164 | 10,371,693 |
|  | TR-1 (bp) | 1,824,356 | 4,984,826 | 4,979,336 |
|  | CentC (bp) | 708,574 | 3,233,759 | 2,939,574 |
|  | rDNA intergenic spacer (bp) | 185,513 | 789,045 | 622,630 |
|  | Subtelomere (bp) | 99,218 | 554,866 | 598,330 |

**Supplementary Table 5. Composition of CentC arrays.**

| Chr | Start (bp) | End (bp) | Size (bp) | 100N^a^ | CentC (%) | CRM (%) | cinful-zeon (%) | grande (%) | huck (%) | opie-ji (%) | prem1 (%) |
| --- | --- | --- | --- | --- | --- | --- | --- | --- | --- | --- | --- |
| chr1 | 133,797,033 | 134,161,088 | 364,055 | N | 28.2 | 1.9 | 9.0 | 3.8 | 6.2 | 2.6 | 3.7 |
| chr1 | 136,702,219 | 138,370,323 | 1,668,104 | Y | 45.7 | 38.8 | 2.4 | 0.0 | 1.5 | 0.9 | 0.6 |
| chr2 | 95,886,484 | 96,018,046 | 131,562 | N | 19.1 | 45.8 | 10.7 | 0.0 | 0.0 | 6.8 | 6.5 |
| chr3 | 83,463,110 | 83,579,253 | 116,143 | N | 29.9 | 0.0 | 6.4 | 12.3 | 11.2 | 0.0 | 6.8 |
| chr3 | 85,837,492 | 86,608,170 | 770,678 | N | 37.8 | 41.3 | 3.8 | 0.0 | 3.7 | 1.1 | 3.3 |
| chr4 | 109,241,724 | 109,363,707 | 121,983 | N | 14.2 | 61.3 | 4.0 | 0.0 | 0.0 | 0.0 | 0.0 |
| chr5 | 107,727,923 | 107,946,850 | 218,927 | N | 41.8 | 34.3 | 4.5 | 0.0 | 0.0 | 0.0 | 3.1 |
| chr6 | 53,802,144 | 54,696,183 | 894,039 | Y | 50.5 | 40.8 | 0.7 | 0.0 | 1.4 | 0.2 | 0.0 |
| chr7 | 56,131,392 | 56,883,231 | 751,840 | Y | 32.6 | 54.5 | 0.3 | 0.0 | 0.0 | 0.0 | 0.0 |
| chr7 | 58,725,452 | 59,026,217 | 300,765 | N | 73.7 | 2.3 | 0.0 | 0.0 | 0.0 | 6.1 | 5.3 |
| chr8 | 49,443,032 | 49,660,040 | 217,008 | N | 60.9 | 23.8 | 3.4 | 0.0 | 0.0 | 0.0 | 3.7 |
| chr8 | 50,101,170 | 50,244,111 | 142,941 | N | 47.8 | 28.6 | 9.9 | 0.0 | 0.0 | 0.0 | 0.0 |
| chr9 | 58,430,781 | 58,672,011 | 241,230 | N | 59.7 | 15.8 | 0.0 | 0.0 | 0.0 | 0.0 | 8.5 |
| chr9 | 58,867,400 | 59002849 | 135,449 | N | 61.2 | 31.0 | 0.4 | 0.0 | 0.0 | 0.0 | 0.0 |
| chr10 | 49,448,433 | 49,559,148 | 110,715 | N | 67.8 | 0.0 | 0.0 | 0.0 | 11.1 | 0.0 | 8.6 |
| chr10 | 52,128,660 | 52,443,218 | 314,558 | N | 49.2 | 31.1 | 5.2 | 0.0 | 0.0 | 4.2 | 3.2 |

^a^ Gaps marked by 100 Ns are of unknown size.

**Supplementary Table 6. Composition of knob180 and TR-1 knobs.**

|  | Chr | Start (bp) | End (bp) | Size (bp) | 100N^a^ | Ngap (%) | knob180 (%) | TR-1 (%) | cinful-zeon (%) | grande (%) | huck (%) | opie-ji (%) | prem1 (%) |
| --- | --- | --- | --- | --- | --- | --- | --- | --- | --- | --- | --- | --- | --- |
| TR-1 | chr4 | 231,661,938 | 232,984,750 | 1,322,813 | N | 0.0 | 0.1 | 51.0 | 30.3 | 0.0 | 0.0 | 1.9 | 1.9 |
|  | chr10 | 142,321,698 | 146,554,927 | 4,233,230 | N | 0.0 | 1.1 | 42.7 | 27.3 | 0.7 | 1.2 | 1.2 | 3.1 |
|  | chr10 | 150,506,817 | 153,088,305 | 2,581,489 | N | 0.0 | 0.0 | 42.0 | 29.7 | 1.2 | 1.7 | 1.4 | 2.2 |
|  | chr10 | 157,208,255 | 159,276,069 | 2,067,815 | N | 0.0 | 0.5 | 36.9 | 20.9 | 0.6 | 2.5 | 4.9 | 5.9 |
| knob180 | chr5 | 198,844,996 | 200,124,258 | 1,279,263 | N | 0.0 | 59.3 | 5.0 | 12.0 | 1.1 | 1.1 | 2.0 | 5.0 |
|  | chr6 | 1 | 623,495 | 623,495 | N | 14.9 | 56.7 | 2.3 | 4.7 | 2.3 | 1.9 | 0.0 | 4.5 |
|  | chr6 | 176,451,347 | 177,079,220 | 627,874 | N | 0.0 | 28.9 | 18.4 | 11.9 | 4.5 | 1.5 | 4.0 | 0.8 |
|  | chr7 | 155,073,718 | 157,644,910 | 2,571,193 | Y | 7.0 | 66.0 | 2.7 | 5.9 | 0.0 | 0.6 | 2.8 | 1.6 |
|  | chr8 | 160,828,702 | 162,735,401 | 1,906,700 | Y | 15.0 | 58.2 | 3.7 | 6.3 | 0.0 | 0.7 | 1.4 | 1.6 |
|  | chr9 | 1,630 | 842,889 | 841,260 | N | 0.0 | 67.0 | 4.3 | 7.4 | 1.7 | 5.9 | 4.3 | 0.9 |
|  | chr10 | 174,217,005 | 178,146,998 | 3,929,994 | Y | 7.8 | 63.6 | 0.8 | 7.6 | 0.4 | 1.7 | 2.0 | 1.5 |
|  | chr10 | 180,357,409 | 182,945,132 | 2,587,724 | N | 0.0 | 53.9 | 0.0 | 9.8 | 1.1 | 0.5 | 5.6 | 4.0 |

^a^ Gaps marked by 100 Ns are of unknown size.

**Supplementary Table 7. Gene and transposon distributions in the Ab10 haplotype and corresponding N10 regions.**

| Region | N10/Ab10 Proximal ^a^ | Ab10 Shared ^b^ | Ab10 Specific ^c^ |
| --- | --- | --- | --- |
| Size (bp) | 20,000,000 | 12,716,384 | 22,438,721 |
| Genes | 521 | 580 | 450 |
| Gene density (genes/Mb) | 26.1 | 45.6 | 20.1 |
| CDS content (%) | 8.8 | 15.0 | 5.5 |
| Average gene length (bp) | 3,377 | 3,284 | 2,757 |
| Average CDS length (bp) | 1,465 | 1,425 | 911 |
| Single exon gene (%) | 35.5 | 36.0 | 49.1 |
| Genes overlapped with TE by 95% (%) | 4.2 | 4.3 | 7.9 |
| TE content (%) | 74.2 | 76.4 | 88.1 |

^a^ Sequence in a 20 Mb region left of the first TR-1 knob that is not a part of the Ab10 haplotype (122.3 - 142.3 Mb).

^b^ Sequence in two large inversions with shared synteny between the Ab10 haplotype and N10 (153.0 -157.4 Mb and 159.4 - 167.7 Mb).

^c^ Sequence present in the Ab10 haplotype but not the B73 N10 genome, including a region between the first two TR-1 knobs (146.6 - 150.5 Mb), a region from the end of the second inversion to the large knob (167.7 - 174.3 Mb), and a region from the large knob to the end of Ab10 haplotype (183.1 - 195.0 Mb).
